# Fibre type differences in the organisation of mononuclear cells and myonuclei at the tips of human myofibres

**DOI:** 10.1101/2024.05.03.592365

**Authors:** Christian Hoegsbjerg, Ask Møbjerg, Ching-Yan Chloé Yeung, Peter Schjerling, Michael R. Krogsgaard, Manuel Koch, Michael Kjaer, Arvind G. von Keudell, Abigail L. Mackey

## Abstract

The myotendinous junction (MTJ) is a weak link in the musculoskeletal system. Here, we isolated the tips of single myofibres from healthy human hamstring muscles for confocal microscopy (n=6) and RNAscope *in situ* hybridization (n=6) to gain insight into the profiles of cells and myonuclei in this region. A marked presence of mononuclear cells was observed coating the fibre tips, with a median of 29 (range 16-63) and 16 (9-23) cells per fibre for type I and II myofibres, respectively (p<0.05). The number and density of myonuclei gradually increased from the myofibre proper towards the tip (p<0.05), similarly for both fibre types, and a greater number of *COL22A1*-expressing nuclei was seen in type II vs type I myofibres (p<0.05). These divergent fibre type-specific characteristics of the MTJ reflect the respective demands for remodelling of the tendon and myofibre sides of the junction according to loading patterns. This insight refines our fundamental understanding of the human MTJ at the cell and structural levels.

**Summary statement:** At the site of attachment to tendon, type I and II human myofibre tips display divergent numbers of mononuclear cells and COL22A1+ nuclei, changing our understanding of myotendinous junction biology.

## Introduction

The proper function of the muscle-tendon-bone unit is paramount for human locomotion and breathing, but the unit is only as strong as its weakest link. The myotendinous junction (MTJ) is the predominant site of injuries in the hamstring muscles, accounting for half of all sports injuries (Edouard et al., 2023; Grange et al., 2023). Susceptibility of the MTJ to injury seems paradoxical given the high degree of structural and molecular specialisation of the MTJ. For example, the muscle-tendon interface interdigitates to enhance contact area (Huijing, 1999; Knudsen et al., 2015; Kvist et al., 1991; Tidball, 1991), with fibre type differences. Type I muscle fibres have a larger interface area than type II fibres at the human (Jakobsen et al., 2023) and frog (Tidball and Daniel, 1986) MTJ, in line with the observation that muscles with a high proportion of type II muscle fibres are more susceptible to injury at the MTJ (Lievens et al., 2022).

In addition to structural specialisation, the MTJ is distinct in its protein composition when compared to neighbouring muscle and tendon tissue (Jacobson et al., 2020; Karlsen et al., 2022). One of the proteins unique to the MTJ in the muscle-tendon unit is collagen XXII (Koch et al., 2004). *COL22A1*, the gene encoding collagen alpha 1 (XXII) chain, is essential for the stability of the interface (Charvet et al., 2013; Malbouyres et al., 2022) and has been detected in both tenocyte-like cells and myonuclei at the MTJ by RNA sequencing at the level of the single cell (Scott et al., 2019; Yan et al., 2022; Yaseen et al., 2021) or single nucleus (Dos Santos et al., 2020; Karlsen et al., 2023; Petrany et al., 2020; Wen et al., 2021). However, since these analyses are performed on tissue homogenate, we lack the spatial dimension and evaluation at the level of the myofibre, the primary unit of attachment of muscle to tendon. Furthermore, due to the disparity in motor unit recruitment patterns and forces exerted between fibre types, it is possible that type I and II fibres have different needs for maintenance and repair, inside and outside the fibre. Interestingly, in an earlier study, we detected markedly fewer *COL22A1*-expressing type I vs type II myonuclei by snRNA-seq in human MTJ tissue (Karlsen et al., 2023), despite a larger surface area of type I myofibre tips (Jakobsen et al., 2023). Thus, many research questions related to the function of the MTJ remain unresolved.

It is assumed that the structural and molecular properties increase the contact area and the strength of protein binding between the myofibres and tendon to maximize the strength of the MTJ and its resistance to strain. However, maintenance and repair require cellular activity on both sides of the junction, raising the question of how strength and maintenance of the MTJ are prioritised, and whether type I and II fibres follow the same strategy. The presence of mononuclear cells can occupy sites for contact between the two tissues, while a high density of myonuclei may interfere with force transmission by disturbing myofilament alignment and attachment to the myofibre tip. Therefore, insight into the arrangement of cells and myonuclei at the MTJ, and its relation to fibre type, is fundamental for our understanding of tissue architecture and dynamic function of this region. To address this, we performed RNAscope and 3D microscopy of the tips of type I and II human muscle fibres, revealing a surprisingly high number of mononuclear cells and *COL22A1*- expressing nuclei, with differences between fibre types. These findings expand the framework for understanding the potential of the MTJ to resist force and to respond to day-to-day loading and severe disruption with injury.

## Results

### Nuclear aggregation at the MTJ at tissue and single myofibre levels

We first investigated the nuclear organisation of the MTJ in early postnatal and adult healthy tissues by immunofluorescence using the cytoskeletal marker vimentin. In *biceps brachii* muscle-tendon tissue from 19-day-old pigs (born 10 days pre-term) we observed a band between the muscle and tendon, strongly stained by vimentin and enriched in nuclei (Fig. 1A,B), also evident in adult human *gracilis* tissue (Fig. 1C). Next, we investigated this phenomenon at the single myofiber level, using high-resolution laser scanning confocal microscopy of fixed single myofibers isolated from the *gracilis* of two patients. The structure of the myofibre-tendon attachment was preserved with the single myofibre isolation procedure, evident from undisrupted folding of the dystrophin-stained sarcolemma adjacent to the collagen XXII-stained basement membrane (Koch et al., 2004), firmly attached to a tail of tendon tissue (Fig. 1D,E, arrows). We observed a clear aggregation of nuclei at the folded region of the myofibre tip (Fig. 1D,E; Movie S1). As the collagen XXII staining closely followed the folded sarcolemma, the length of the sarcolemmal foldings was used to define the MTJ region in the subsequent analyses (Fig. 1D,E).

**Figure 1.**
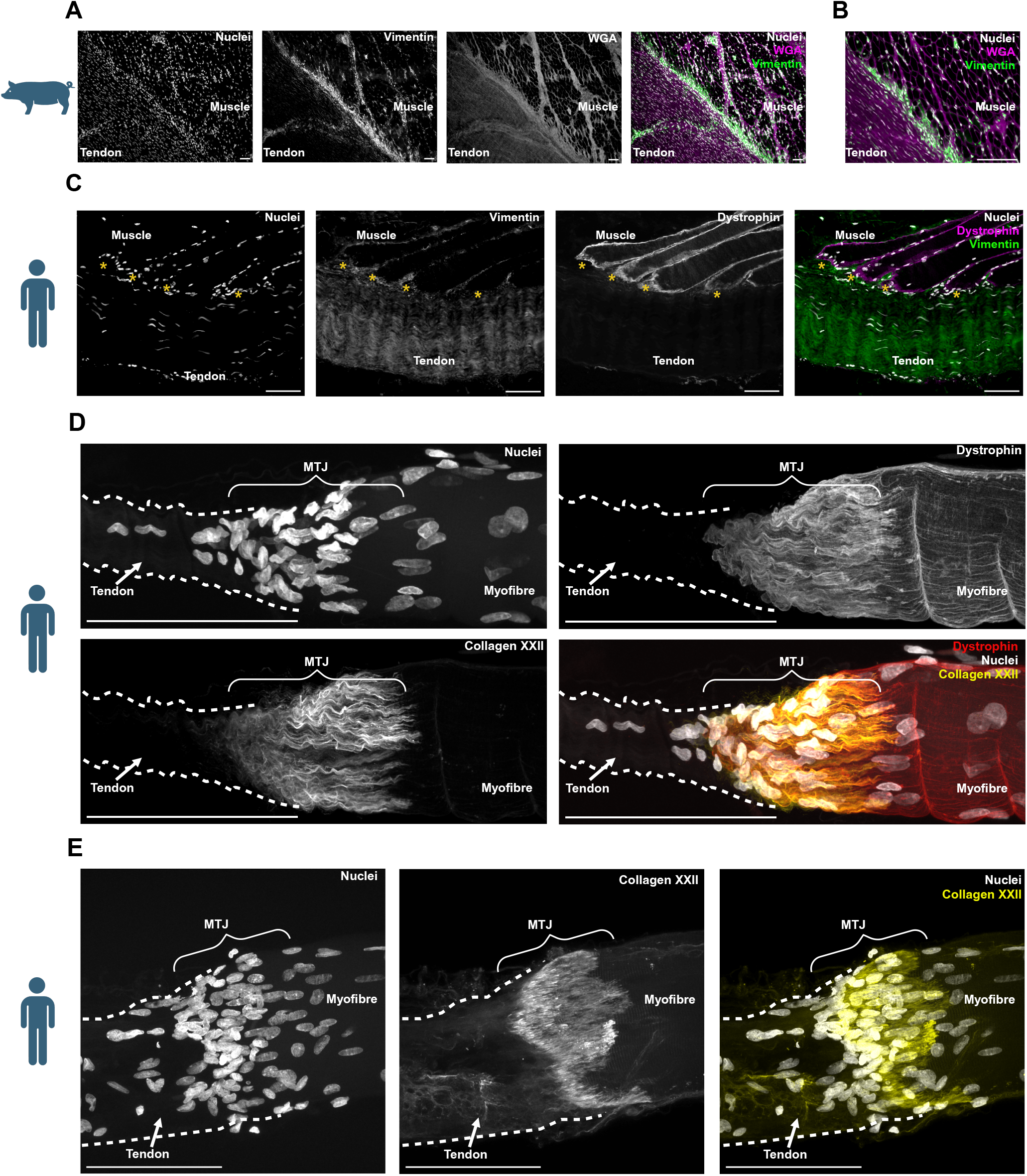
Immunofluorescence of the MTJ. Scale bars: 100 µm. Images are representative of n=2 patients/pigs. **A:** Single channel greyscale and composite images of widefield immunofluorescence microscopy of 10 µm thick sections of pre-term pig *extensor carpi radialis* muscle-tendon tissue, stained for vimentin, wheat germ agglutinin (WGA), and nuclei. A large band of vimentin+ cells is present between the muscle and tendon tissue. **B:** Upper left part of section in A acquired with higher magnification. **C:** Single channel greyscale and composite images of widefield immunofluorescence microscopy of 10 µm thick sections of longitudinally cut human *gracilis* muscle-tendon tissue. Sections stained for vimentin, dystrophin and nuclei. Many nuclei are visible in the muscle-tendon interface. *insertion of myofibres into the tendon. **D,E:** Maximum Z-projection single channel greyscale and composite images of laser scanning confocal microscopy of human *gracilis* single myofiber fragments teased off the tendon. The fibres have been stained for collagen XXII, dystrophin and nuclei (D), or collagen XXII and nuclei (E). A large aggregation of nuclei can be seen at the MTJ of both myofibers.

### 3D model of the spatial arrangement of nuclei at the MTJ

To investigate the proportion of nuclei (observed in Fig. 1D,E) that were either nuclei of mononuclear cells or myonuclei, we utilised a 3D approach. From confocal images of 10 type I and 10 type II myofibres from the *gracilis* muscle of each patient (n = 6, no reported history of disease), a computer model was generated of the myofibre actin core (Fig. 2A). Each nucleus was manually evaluated and classified as either being a nucleus outside the myofibre (dystrophin-defined sarcolemma) or a myonucleus (inside the dystrophin-defined sarcolemma, and vimentin negative). For outside cells, we used a cut-off distance from the myofibre surface of 10 µm as this is the approximate diameter of a fibroblast (Uhal et al., 1998) and these cells are theoretically close enough to the myofibre for cell-cell contact (Fig. 2A,B). We classified these cells as MTJ cells. The majority of the MTJ cell nuclei were within 2-3 µm of the myofibre surface (Fig. S1A-C). All data calculated for the individual myofibres included in the subsequent figures can be found in the supplemental data S1.

**Figure 2.**
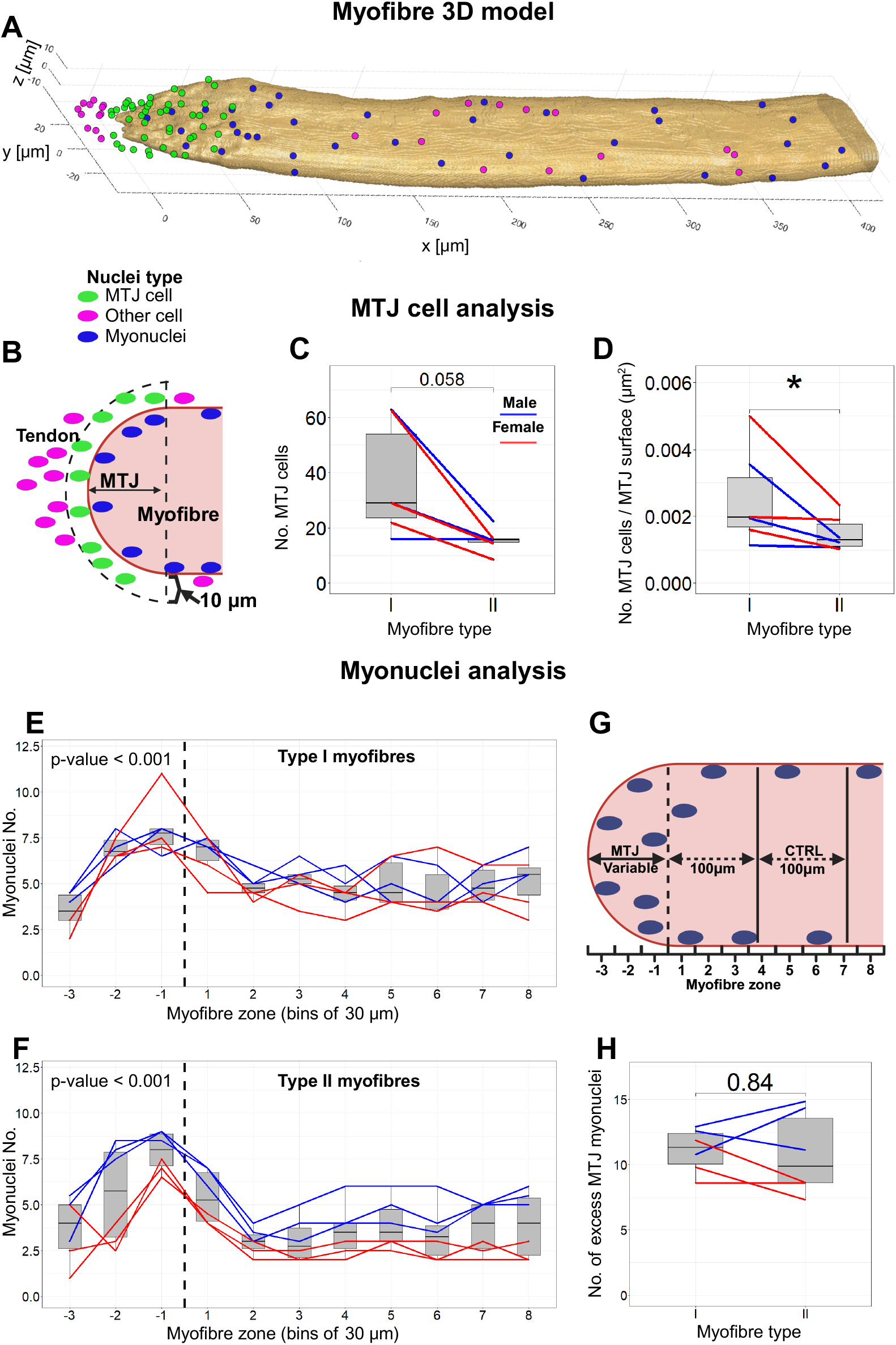
3D model quantification of nuclei. From spinning disk images on single human myofibre tips (see Figure 6.C), a 3D model was generated to quantify nuclei inside and outside the myofibre. Individual patient medians are displayed; n=6 patients. Boxplots and whiskers are calculated according to the Tukey method. Created using Biorender.com. **A:** Example of 3D model output from matlab. The model is based on actin staining and includes the surface rendering of the myofibre with the nuclei spots overlayed. Blue spots are myonuclei, green spots are nuclei of MTJ cells, and magenta spots are nuclei of excluded outside cells. **B:** 2D schematic of the MTJ cell 3D analysis, quantified in C and D. Mononuclear cells were included as MTJ cells if the nuclei spot was located within 10 µm of the myofibre MTJ surface. Image not to scale. **C,D:** Quantification of MTJ cells at the tips of type I and II myofibres. Displayed are both absolute numbers per myofibre (C) and number of MTJ cells normalized to the MTJ surface area (D). * p < 0.05 between fibre types, Wilcoxon signed rank test. **E,F:** Profiles of myonuclear numbers in type I and II myofibres. The dotted line is the MTJ border depicted in G. Zones are arranged in increments of 30 µm from the MTJ border as indicated in G. Analysed by a Friedman rank sum test. **G:** 2D schematic of MTJ myonuclei 3D analysis. Per myofibre, myonuclei were classified as MTJ if they were between the myofibre tip and the MTJ border, or as belonging to the control region (CTRL) if they were within a 100 µm bracket displaced by 100 µm from the MTJ. For myonuclear number profiles (E, F), myonuclei were allocated bins of 30 µm increments, so that bin (zone) 1 and -1 is the first zone towards the myofibre proper and myofibre tip, respectively. Dotted line indicates MTJ border. Drawing not to scale. **H:** Number of excess myonuclei at the MTJ (the expected number (CTRL^myonuclei^ ^no.^ * (MTJ^volume^/CTRL^volume^)) subtracted from the observed number). MTJ and CTRL myonuclei are defined in G. Data are analysed by Wilcoxon signed rank test.

### Marked presence of cells coating myofibre tips and fibre type differences

Each myofibre tip was associated with a surprisingly large population of MTJ cells, with a median of 29 (range 16-63) MTJ cells per type I myofibre and 16 (9-23) cells per type II myofibre (Fig. 2C). Type I myofibres displayed larger variation between subjects than type II myofibres. Nevertheless, there was a clear trend for a larger MTJ cell pool at the tips of type I myofibres compared to type II (p = 0.058), which was statistically significant when normalized to the MTJ surface area (p = 0.031) (Fig. 2C,D). Notably, the MTJ surface area was larger for type I myofibres (35% larger, p-value = 0.031, Fig. S1D), as previously reported with a similar approach (Jakobsen et al., 2023).

### Elevated number of myonuclei at myofibre tips, independent of fibre type

Next, we used the model to investigate the number of myonuclei at the tips of single myofibers to determine if they also contribute to the overall aggregation of nuclei at the MTJ. When analysing the profile of myonuclear numbers at the tip and myofibre proper, we found both fibre types display an elevated number of myonuclei at the fibre tip (p<0.001, Friedman test) with a similar peak in numbers (zone -1, Fig. 2E,F). To further investigate this elevation in myonuclear numbers, we calculated the predicted number of myonuclei at the tip (MTJ) based on the relationship between myonuclear content and myofibre volume in a control (CTRL) segment, which was 100 µm in length and separated by the MTJ by an additional 100 µm along the myofibre (Fig. 2G). The tips of both fibre types displayed an excess of ∼11 myonuclei (ranges of excess myonuclei were 9-13 and 7-15 for type I and II fibres, respectively, Fig. 2H) above the predicted number of 9 (range 5-13) for type I and 5 (range 3-11) for type II fibres. (Observed number of MTJ myonuclei were 21 and 16 for type I and II fibres, respectively, Fig. S1E). Additional measurements for the MTJ and CTRL regions can be found in Fig. S1.

#### Altered spatial organization of myonuclei approaching the MTJ

To further explore the higher number of myonuclei at the MTJ, we used the model to investigate changes in the organization of myonuclei at the myofibre tips compared to the myofibre proper. We calculated the size of the myonuclear domain (MND), which is the volume of cytoplasm that can be attributed to an individual myonucleus (Fig. 3A). As with the number of excess myonuclei, we observed no difference in the MND size at the MTJ between fibre types (Fig. 3B). Both fibre types displayed the same pattern of a markedly lower MND size at the fibre tip, which then increased in size along the fibre before reaching a plateau (Fig. 3C,D). A similar profile was observed for nearest neighbour (NN), a measurement of myonuclear proximity and a proxy for density (Fig. 3E-G). A difference in NN at the MTJ was observed between fibre types (Fig. S1I). No difference in MND size or NN was observed between fibre types in the CTRL region, although trends for a lower number of myonuclei per 100 µm and a larger MND size were found in type II myofibres (Fig. S1J- M), which is in line with previous reports (Cristea et al., 2010). When comparing the MND size between the CTRL and MTJ regions we found that both fibre types showed significantly smaller domain sizes at the MTJ. Type I myofibres displayed a 2.7-fold difference (12334 µm^3^ vs. 4508 µm^3^, p < 0.05, Fig. 3I) and type II a 3.4-fold difference (13360 µm^3^ vs. 3929 µm^3^, p < 0.05, Fig. 3J). The same was true for the difference in NN between CTRL and MTJ regions for both fibre types, although less pronounced with a 1.6- and 1.9-fold difference in type I and II myofibres, respectively (p < 0.05, Fig. 3K,L).

**Figure 3.**
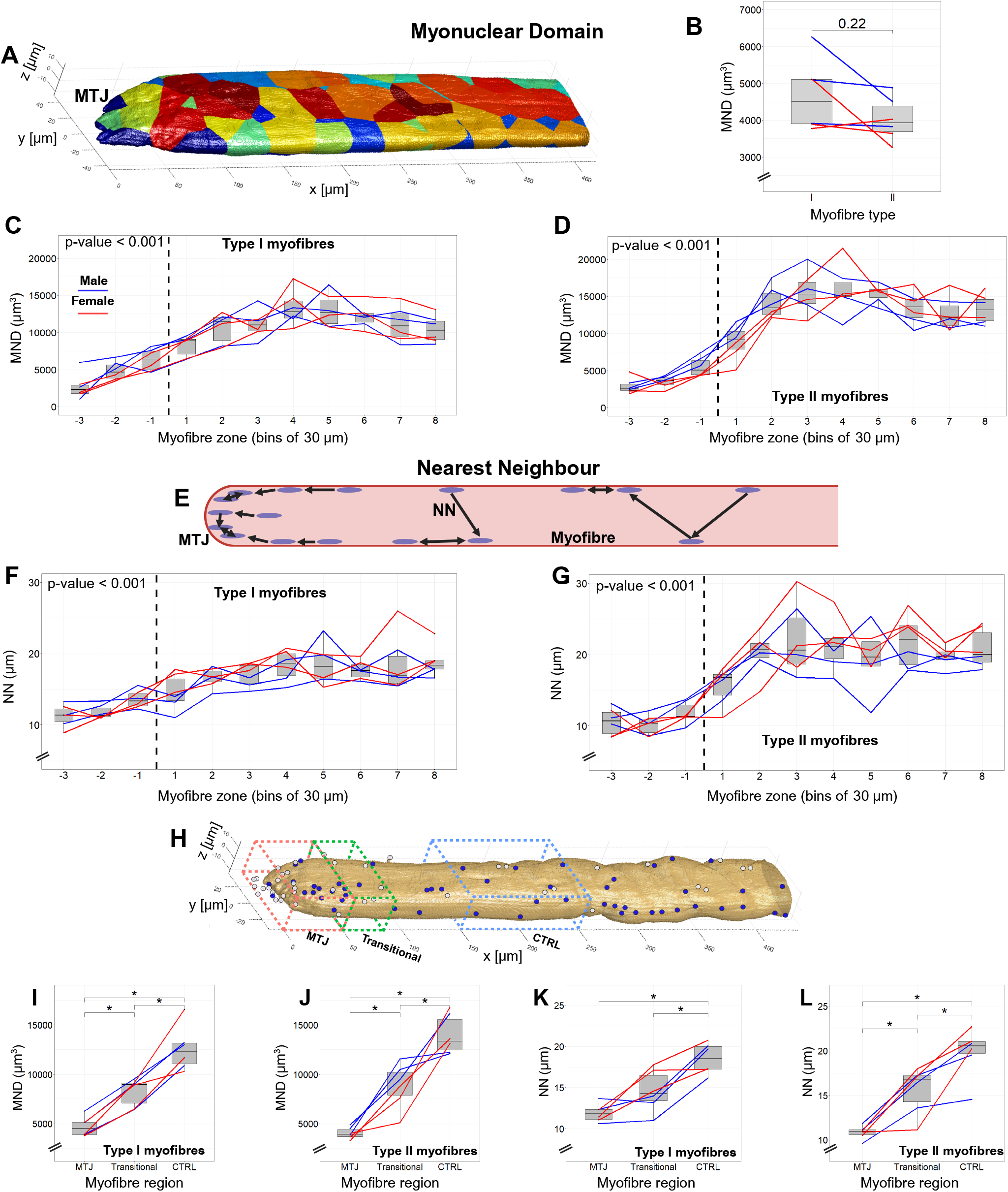
– Spatial organization of myonuclei at the MTJ. Given the strong relationship between myonuclear content and myofibre volume, we determined MyoNuclear Domain (MND) size and Nearest Neighbour (NN) distance. Individual patient medians are displayed; n=6 patients. Boxplots and whiskers are calculated according to the Tukey method. Created using Biorender.com. **A:** Matlab image example of calculated myonuclear domains (MND). The leftmost voxel of the myofibre surface (MTJ) is used to index the x-axis at position 0. **B:** MTJ MND size in type I and II myofibres. Analysed by Wilcoxon signed rank test. **C-D:** MND profiles of type I and II myofibres, respectively. The profile is derived from the same zones as in Fig. 2E,F,G. Analyzed by Friedman rank sum test. **E:** Schematic of NN calculation. Each myonucleus is assigned a distance value in µm to the nearest other myonuclei. The schematic is not to scale. **F-G:** As C-D but showing the NN profile. **H:** Matlab output image showing myofibre and nuclei segmentation. The regions analysed in I-L are indicated. MTJ (red box) and CTRL (blue box) region is the same as 2E. In I-L we included a transitional region (green) which is a fixed region always extending 30 µm immediately after the MTJ region. Blue and white dots are myonuclei and nuclei outside the myofibre, respectively. **I-L:** Boxplots MND and NN for the regions depicted in H. * p < 0.05 between regions, Wilcoxon signed rank test.

Interestingly, for both fibre types, the plateau in MND size and NN is reached approximately 30 µm after the end of the MTJ region (zone 1, Fig. 2, 3). This 30 µm region may represent the transitional region between the MTJ and main body of the myofibre we proposed recently (Fig. 3H-L) (Karlsen et al., 2023).

### A greater number of *COL22A1*-expressing nuclei at the MTJ of type II vs type I myofibres

To follow up on our previous observation by snRNA-seq of an unexpectedly low number of *COL22A1*-expressing myonuclei in type I fibres (Karlsen et al., 2023), we used RNAscope to quantify *COL22A1*-expressing nuclei from Z-stack maximum intensity projections of type I and II single myofibres from the *semitendinosus* muscle of 6 patients (Fig. 4A). In accordance with the snRNA-seq data of the tenocyte cluster (Fig. S3A,B), some nuclei outside the myofibre were positive for *COL22A1* transcripts (Fig. 4A, arrow), but the majority of *COL22A1*-positive nuclei were within the myofibre. Type II myofibres displayed more *COL22A1*-positive nuclei than type I (median 16 (range 4-53) vs 9 (range 3-21), p = 0.036, Fig. 4B). Consistently, for all fibres across all patients, *COL22A1* was localised to the tip region of the fibre, as expected from immunofluorescence staining (Fig. 1D,E) and snRNA-seq data (Fig. S3). Furthermore, the distribution of *COL22A1*-positive nuclei (Fig. 4C,D) mirrors that of the absolute number of myonuclei at the MTJ (Fig. 2E,F).

**Figure 4.**
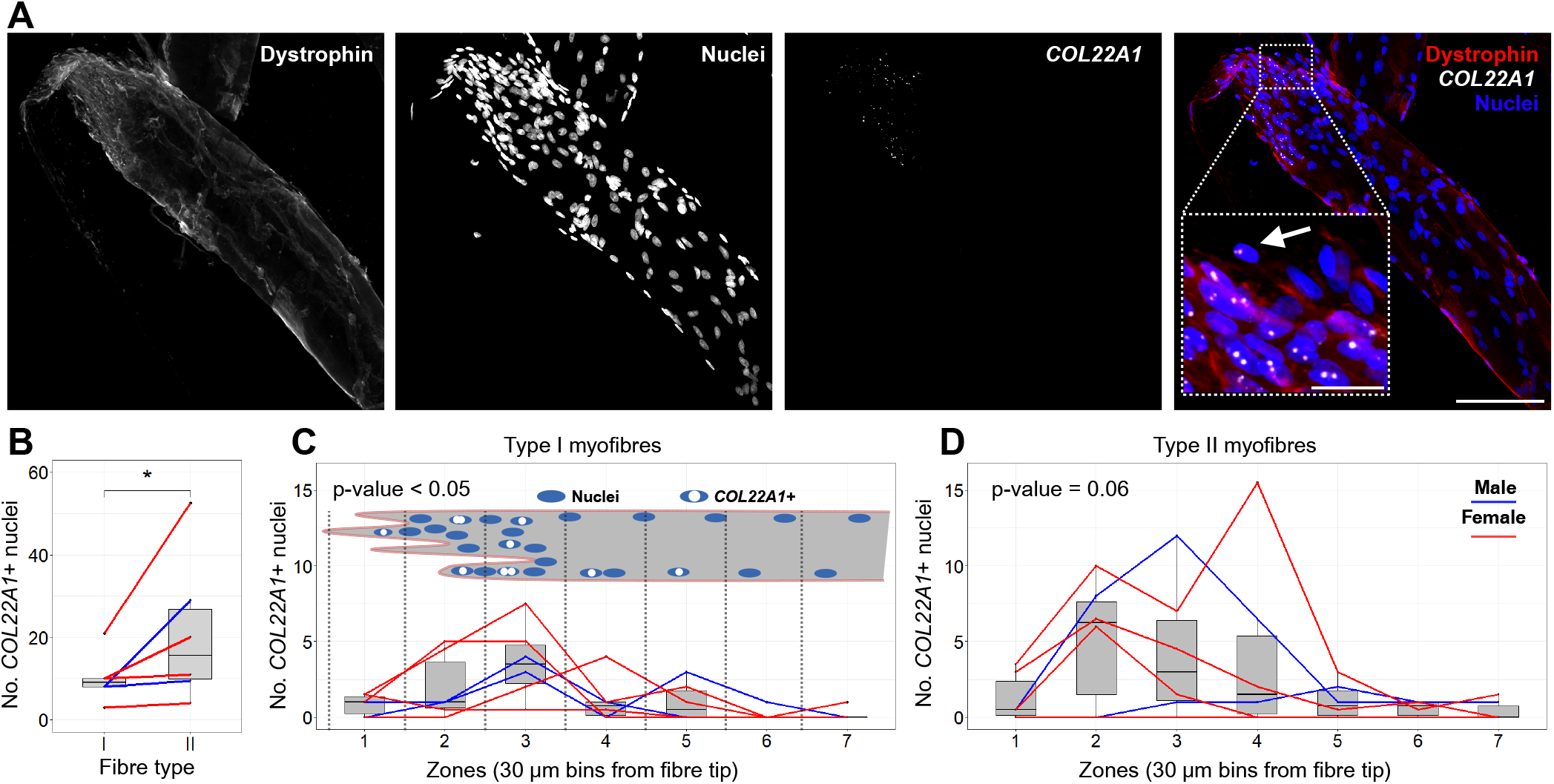
MTJ nuclear *COL22A1* expression. RNAscope of fixed human single myofibre tips. Individual patient medians are displayed; n=6 patients. Boxplots and whiskers are calculated according to the Tukey method. Created using Biorender.com. **A**: Example of Z-stack maximum intensity projection of a laser scanning confocal image showing a single myofibre stained for dystrophin, nuclei and an RNAscope probe targeting *COL22A1* RNA. **B**: Boxplots showing the number of *COL22A1*+ nuclei. *P < 0.05, Wilcoxon singed rank test. **C,D**: Distribution of *COL22A1*+ nuclei. Boxplots show number of *COL22A1*+ nuclei in increments of 30 µm from the myofibre tip (as displayed in inlet of C) in type I (C) and II (D) myofibres. Analysed by a Friedman rank sum test.

## Discussion

Here we report muscle fibre type dependent and independent profiles of cells and myonuclei at the tips of human myofibres from healthy individuals. Myofibre tips were characterised by a high density of myonuclei and a high number of mononuclear cells closely associated with the fibre surface. Divergent fibre type differences were observed with a higher number of mononuclear cells on type I compared to type II myofibres, while, in contrast, type II myofibre tips displayed a higher number of *COL22A1*-expressing nuclei. These findings are in line with higher requirements for maintenance and repair of the myofibre basement membrane of type II fibres and the tendon site of attachment of type I fibres, according to fibre type specific patterns of muscle fibre recruitment and time under tension.

It is generally assumed that the increased surface area of the MTJ reduces stress during the propagation of force from muscle to tendon (Charvet et al., 2012; Jakobsen and Krogsgaard, 2021; Tidball, 1991; Tong et al., 2024). Counterintuitively, we find a large presence of mononuclear cells closely associated with the myofibre tip, exceeding 60 cells per fibre for some patients, and in theory taking up space for force dissipation. This finding may indicate a lesser importance of the MTJ in force transmission, and instead lend further support for the role of lateral force transmission (Huijing, 1999). However, it is also possible these cells contribute directly to force transmission by bridging the myofibres and tendon, as previously proposed (Yan et al., 2022). Fibroblasts have been seen by electron microscopy extending their cytoplasm into tendon interdigitations at the MTJ of three-week old rat muscle, with distances of only 50nm between the plasmalemma and the sarcolemma (Mackay et al., 1969). However, no evidence exists for the presence of direct myofibre- cell attachments or force transmission. An alternative role is the maintenance and repair of the MTJ. Our RNAscope data show that *COL22A1* is expressed by MTJ cells (as well as MTJ myonuclei) in human adult muscle, in line with other studies (Dos Santos et al., 2020; Karlsen et al., 2023; Petrany et al., 2020; Scott et al., 2019; Wen et al., 2021; Yan et al., 2022). The expansion of the *COL22A1^+^* tenocyte population following injury (Scott et al., 2019) favours the idea of a regenerative role for these cells. However, the sheer number of MTJ cells we observed here does not align well with the high susceptibility of the MTJ to injury and the poor regenerative capacity (Grange et al., 2023; Wangensteen et al., 2016), potentially indicating the need for alternative rehabilitation strategies to provide an optimal environment for activity of these cells to improve repair outcomes.

At the myofibre tips, we found an aggregation of myonuclei, along with the localised expression of *COL22A1*. Myonuclear specialisation at the MTJ has been demonstrated by snRNA-seq data (Dos Santos et al., 2020; Karlsen et al., 2023; Kim et al., 2020; Petrany et al., 2020), showing a different transcriptional profile, including expression of *COL22A1*. Here, we not only show the direct localisation of *COL22A1^+^* nuclei to the MTJ (Fig. 4), but also the additional increase in both total number and density of myonuclei at the myofibre tips compared to the main portion of the fibre (Fig. 2E-H, 3). The question then arises regarding the mechanisms driving myonuclear accretion and transcriptional divergence from the myofibre proper. The susceptibility to injury of the MTJ (Grange et al., 2023) indicates greater requirements for maintenance and repair of this region. To meet this demand, the myofibre tips may therefore increase the number of myonuclei, which could be accomplished by a greater rate of myonuclear incorporation, or a higher rate of transport of existing myonuclei to the tips. Myonuclei move towards the MTJ following stretch in drosophila (Perillo and Folker, 2018) and towards sites of focal myofibre damage in mouse muscle (Roman et al., 2021). The elevated need for repair at the MTJ could also explain the higher prevalence of centralised myonuclei, commonly associated with remodelling and myogenesis (Collins et al., 2024; Folker and Baylies, 2013; Jakobsen et al., 2018). Although myonuclear migration to the MTJ is an interesting hypothesis, it does not explain the mechanism governing the altered transcriptional profile of the MTJ myonuclei. One possibility is the origin of the myonuclei. Fusion of fibroblast-like myo-tenogenic (dual identity) cells with the myofibre tip (Esteves de Lima et al., 2021; Yan et al., 2022; Yaseen et al., 2021) could provide new myonuclei with an altered expression profile.

However, this has only been tested in animal developmental and cell culture models, lacking validation in mature tissues, particularly human. It is possible the MTJ cells play a role in the regulation of transcriptional specialisation of myonuclei at the fibre tip, in a similar manner to that observed at the neuromuscular junction (NMJ). Expression of important genes for the NMJ, such as acetylcholine receptors, is restricted to a handful of myonuclei in this region (Denes et al., 2021) and is regulated by the presence of the motoneurone (Anderson and Cohen, 1977; Sanes and Lichtman, 2001). Whether a similar mechanism exists for confining myonuclei, and transcription of MTJ genes, to the myofibre tips remains to be determined.

The distinction between myofibre types is a key feature of our approach and led to the two most notable findings of the study; a higher number of cells associated with the tips of type I myofibres, and a higher number of *COL22A1* expressing nuclei at the tips of type II myofibres. These divergent fibre type adaptations reflect the well-known fibre type differences in myofibre recruitment and force levels. As discussed in detail by James Tidball, it is not only the maximum force reached during a muscle contraction but the time under tension that is important for tissue failure (Tidball, 1991). Type I muscle fibres are recruited more frequently than type II to perform low force movements and to maintain postural control and it is possible that this cyclic, lower loading induces local tendon damage at the attachment sites of type I myofibres. In support of this, molecular damage of tendon collagen I occurs after repeated sub-failure loading with low (6-7%) peak strain (Zitnay et al., 2017). The higher number of cells coating the surface of type I myofibres is in line with a higher requirement for repair of damage to the tendon caused by this loading pattern. Accordingly, the MTJ of type I has a larger surface area than type II myofibres in human (Jakobsen et al., 2023) and frog muscle (Tidball and Daniel, 1986), a finding that is confirmed by our data (Fig. S1F). In contrast, the higher forces exerted by type II myofibres may lead to an alternative adaptation such as the higher number of nuclei expressing *COL22A1* we observed at the fibre tips. This would explain why perturbations in *COL22A1* expression are linked to both the disruption of the MTJ in zebrafish (Charvet et al., 2013; Malbouyres et al., 2022) and an increased risk of muscle strain injury in humans (Miyamoto-Mikami et al., 2020). A deeper understanding of fibre type differences at the MTJ is important for understanding how exercise can protect against injury.

Another interesting finding of our study is the graded specialisation of the MTJ. A change in myonuclear number and organisation started approximately 30 µm before the characteristic sarcolemmal foldings of the MTJ (zone 1 Fig. 2E,F, 3). Our previous snRNA-seq of human MTJ tissue revealed distinct subpopulations of “MTJ” myonuclei, strongly indicating that myonuclei progressively display a more MTJ specific gene expression profile approaching the fibre tip (Karlsen et al., 2023). Taken together with the different expression patterns of MTJ-specific proteins (Jakobsen et al., 2021; Karlsen et al., 2022; Karlsen et al., 2023), we proposed that the transition from myofibre proper to myofibre tip is more graded in nature, rather than a hard border, showing higher degrees of myonuclear specialisation approaching the fibre tips. The tendon-bone enthesis is another example of a graded transition between two tissues of differing mechanical properties (Benjamin et al., 2006; Loukopoulou et al., 2022). The MTJ is thus not merely the interface where muscle meets tendon, but rather several zones: a band of tendon, a large population of mononuclear cells, and a gradient of structural and myonuclear specialisation from the myofibre tips to the myofibre proper. With such a complex architecture, perhaps it is not surprising that regeneration of strain injuries is slow and often incomplete at the structural level (Bayer et al., 2018;

Bayer et al., 2021). Indeed, the poor ability to regenerate the enthesis raises the question of whether it is even possible to recreate the MTJ after separation by strain rupture. Mapping the MTJ zones in detail may help understand why it is disproportionally affected in injuries and in rethinking rehabilitation strategies for complete tissue repair.

In conclusion, our mapping of the spatial profile of nuclei at the muscle-tendon interface at the single myofibre level reshapes understanding of this critical region for force transmission. We further unveil fibre type differences at the MTJ, with a larger pool of MTJ cells residing at the muscle-tendon interface of type I myofibres, while type II myofibres exhibit a higher number of *COL22A1* expressing nuclei. Together these findings are important in understanding the mechanisms governing MTJ cell specialisation and repair, which is essential for improving the clinical outcomes of injuries to the MTJ.

## Materials and methods

### Animals and tissue acquisition

The pig tissue was obtained from a larger study, conducted in accordance with the guidelines from Directive 2010/63/EU of the European Parliament and the study protocol was approved by the Danish Animal Experiments Inspectorate (licence no.: 2020-15-0201-00520), and published elsewhere (Rasmussen et al., 2023). The pigs (Landrace × Yorkshire × Duroc) were born by caesarean section at gestational day 106 (90% of gestation) and reared for 19 days before being euthanised by intracardial sodium-pentobarbital (euthanimal, ScanVet, Animal Health, Denmark). Within 40-60 min, distal samples of the *extensor carpi radialis* containing both muscle and tendon were collected, embedded in TissueTek, frozen in isopentane pre-cooled in liquid nitrogen and stored at -80°C.

### Human patients and tissue acquisition

All human patients of this study provided informed written consent before participation, and the study was approved by the Research Ethics Committee of the Capital Region of Denmark (ref. nr: H-20044907 and ref. nr: H-3-2010-070) and conformed to the Declaration of Helsinki II. Inclusion criteria were that patients were scheduled for ACL reconstruction surgery and had no known diseases or chronic illnesses. 12 patients, 5 males and 7 females (age: 21-40), were included in this study. None performed regular heavy resistance exercise of the hamstring muscles in the month leading up to surgery.

During surgery, both *semitendinosus* (6 patients, 4 females and 2 males) and *gracilis* (6 patients, 3 females and 3 males) waste tendon graft tissue, with muscle tissue still firmly attached, was collected in 50 ml Falcon tubes containing phosphate-buffered saline (PBS, cat. No. P4417- 100TAB, Sigma Aldrich) on ice, within minutes of harvest. While on ice, the tissue samples were divided into smaller pieces. Tissue for cryosections was embedded in Tissue-Tek (Sakura Finetek, Europe, AJ Alphen aan den Rijn, The Netherlands) and frozen in liquid nitrogen-cooled isopentane, before storage at -80 °C. Tissue for the single myofibre immunofluorescence staining, RNA-scope and 3D analysis was fixed by adapting a previous protocol (Mackey and Kjaer, 2017). Briefly, the tissue samples were fixed for 30 minutes at room temperature with Stefanini fixative (Phosphate buffer (0.1mol), containing 2% formaldehyde and 0.15% picric acid, pH 7.2), followed by 4 hours in new Stefanini fixative at 4°C. After fixation, the tissue samples were transferred to 50% glycerol/PBS, left overnight at 4°C, and stored at -20°C. Semitendinosus samples were used for RNAscope in situ hybridisation, and gracilis samples were used for all other techniques.

### Sample preparation and immunofluorescence staining

#### Tissue cryosections

10 µm sections of Tissue-Tek embedded pig or human tissue was cut in either the cross-sectional or longitudinal plane of the muscle fibres with a cryostat at -20°C, collected on glass slides, and stored at -80°C. Before immunofluorescence, slides were moved to room temperature and the sections allowed to dry. After 10 min fixation with Histofix (Cat. no.; 01000, Histolab Products AB, Gothenburg, Sweden), sections were incubated overnight at 4°C with the primary antibodies (diluted in 1% bovine serum albumin (BSA, cat. no.: A3912-50G, Sigma-Aldrich) in tris-buffered saline (TBS; Tris-HCl 0.05 M, sodium chloride 0.154 M, pH 7.4-7.6)), followed by incubation for 45 min at 4°C with secondary antibodies and Hoechst 33342 (unless DAPI was used) diluted in TBS. The samples were washed twice for 10 min in TBS between all protocol steps, and samples were mounted with coverslips and mounting medium with DAPI (Molecular Probes Prolong Gold anti-fade mountant with DAPI, cat. no.: P36931, Invitrogen, Taastrup, Denmark) for pig sections or without DAPI (Molecular Probes Prolong Gold anti-fade mountant, cat. no.: P36930, Invitrogen, Taastrup, Denmark) for human sections. For the full antibody list see Table S2.

#### Single myofibres – Immunofluorescence

Adapting a previous protocol for isolating and staining single muscle fibres (Karlsen et al., 2023; Mackey and Kjaer, 2017), a bundle of myofibres was teased from the tendon of the fixed sample in 50% glycerol/PBS using forceps and a stereo dissection microscope. The bundles were moved to TBS and individual myofibres were teased from the bundle and divided into two fragments, one containing the MTJ with a tendon trail attached and one for myofibre type determination by SERCA staining (Fig. 6A,B, Fig. S4). The MTJ-fragments were then transferred to a 96-well nunc plate and incubated for 1 h at room temperature in immunobuffer (IB: PBS, 50mM glycine, 0.25% BSA, 0.03% saponin, 0.05% sodium azide) with 0.1% Triton X-100 (cat. no. 9036-19-5, Sigma-Aldrich). Primary antibodies, together with Alexa-Fluor 680 Phalloidin, were diluted in IB with 0.1% Triton X-100 and added to the wells for overnight incubation on a rocking table at room temperature, followed by incubation for 2 h with secondary antibodies and Hoechst 33342 diluted in IB (no Triton). The MTJ fragments were aligned in a drop of mounting medium before the coverslip was applied. For full antibody list see table S2.

**Figure 5.**
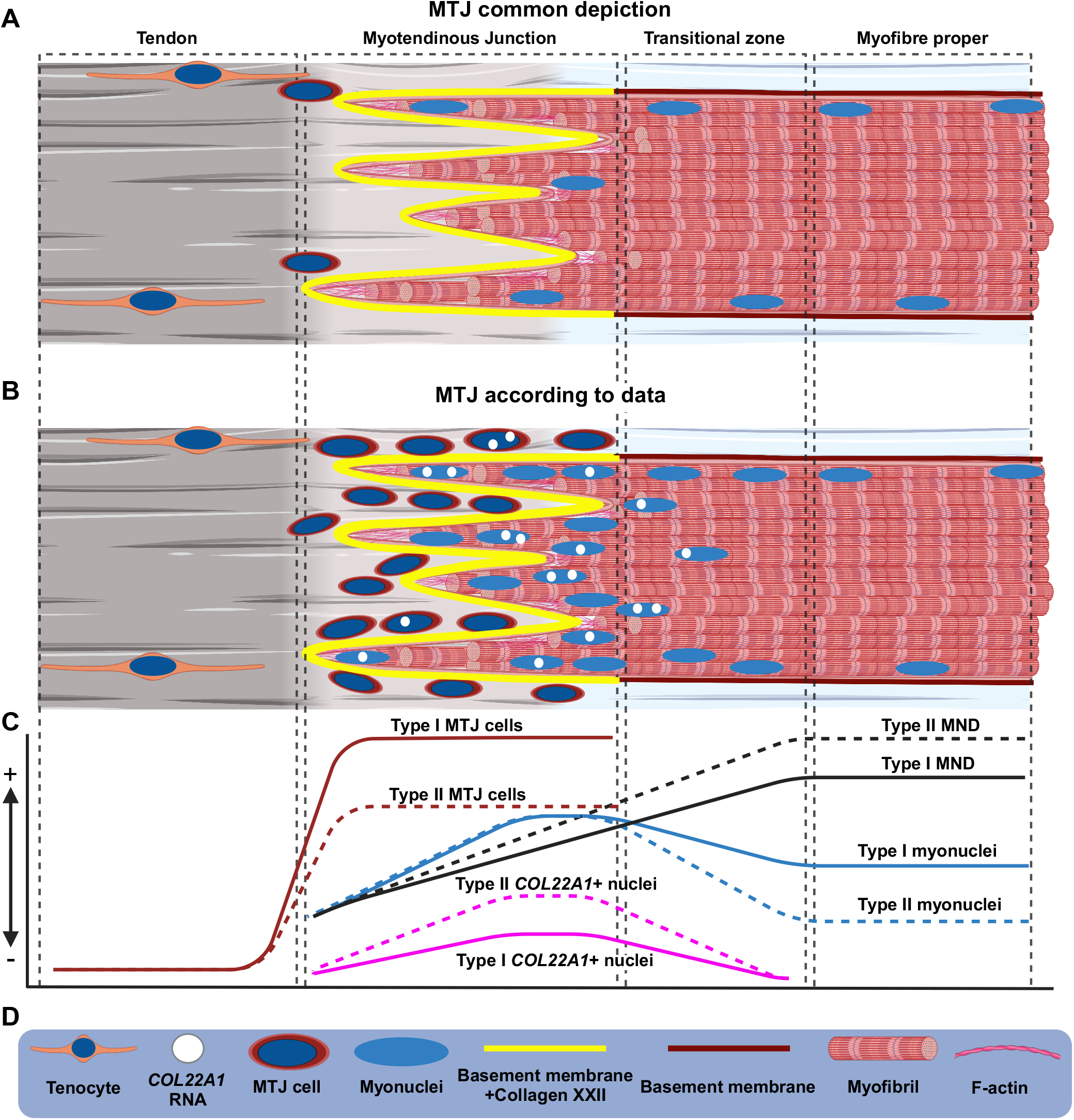
Graphical summary of main findings. Not to scale. Created using Biorender.com. **A:** The general depiction of the myotendinous junction (MTJ). Although the ultrastructure is well characterized, and recent papers suggest both specialized myonuclei and cell populations at the MTJ, the large aggregation of cells and myonuclei at the MTJ are not commonly appreciated. **B:** The MTJ according to our data. On a single myofiber level, the tendon-muscle interface can be separated into distinct zones: ***The tendon***, with low cellular density, ***the MTJ*** with collagen XXII expression, high cellular density, and high myonuclear density, a ***transitional zone*** with steadily decreasing myonuclear number/density, and finally the “***myofibre proper***” with stabilised myonuclear number/density. **C:** A graphical summary of the data presented in this paper showing the relationship between the different zones, type I (solid line) and type II (dashed line) myofibres, the MTJ cell numbers, myonuclear domain (MNDs) sizes, and COL22AI expressing nuclei. **D:** Key symbols used in A and B.

**Figure 6.**
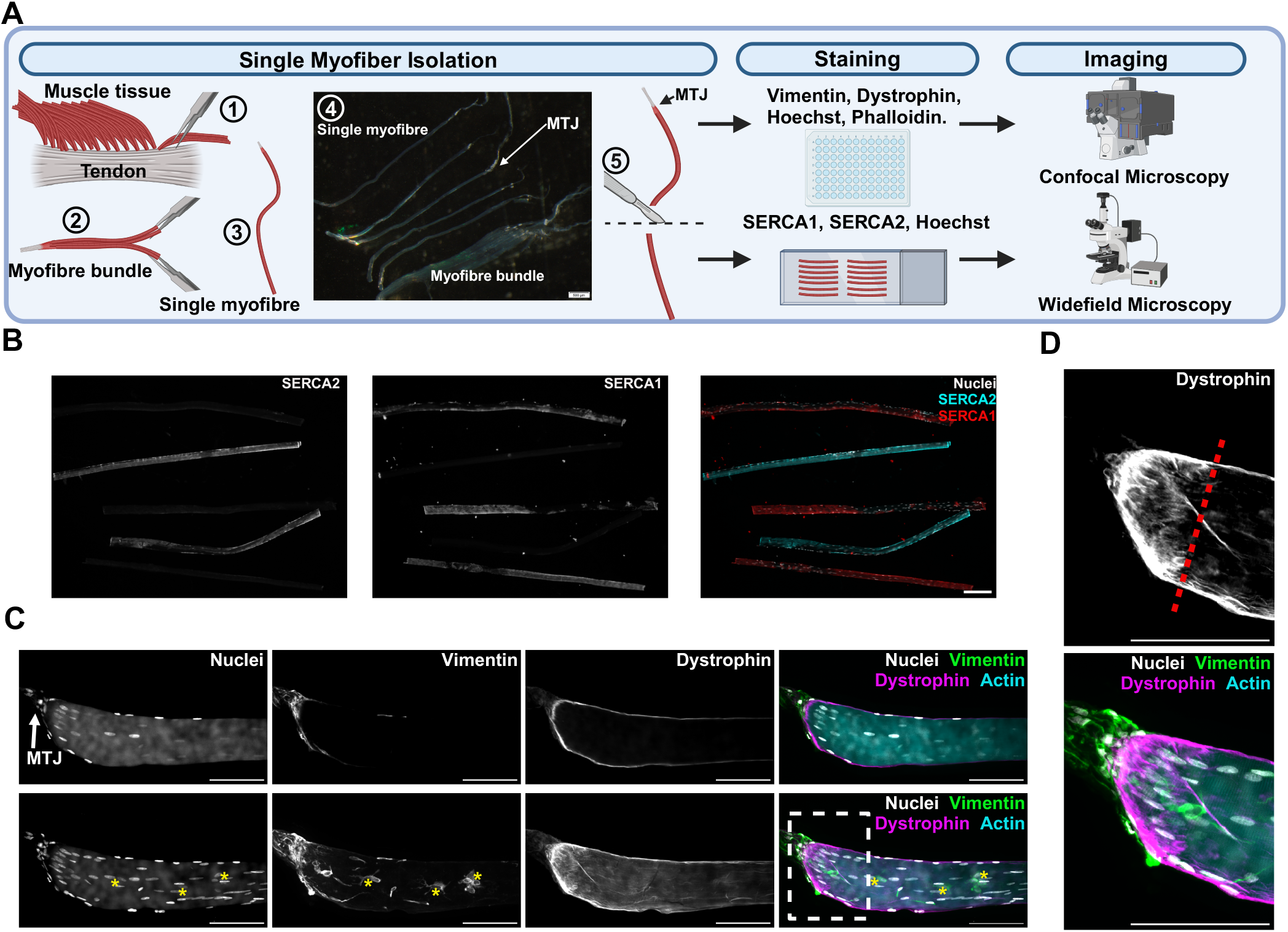
Methodological overview. Scale bars: 100 µm. Created using Biorender.com. **A:** Methodological overview. 1) Bundles of fixed myofibres were teased from the tendon, bringing a portion of tendon with each bundle. 2) Bundles were continuously halved until single myofibres 3) with a tendon tail was obtained. 4) Dissection microscope image of single myofibers (with the MTJ indicated) and visible tendon remnants. 5) The individual myofibres were cut in half, and the MTJ fragment was stained and imaged by confocal microscopy, while the remaining fragment was stained to determine the myofibre type. Full display of final Imaris 3D model generation of MTJ fragments can be viewed in movie S2. **B:** Image example of SERCA staining of type I and II myofibres, indicated by SERCA2 and SERCA1, respectively. **C:** Confocal microscopy image of an MTJ fragment. The upper panel shows a single slice image through the centre of the myofibre, and the lower panel shows a maximum Z-projection of the full stack of images. * highlights perinuclear vimentin staining, which would be classified as nucleus of a mononuclear cell outside the myofibre. Note the high number of myonuclei in the MTJ region. **D:** Magnification of the highlighted region in C. The red dotted line indicates the manually defined MTJ region.

The myofibre typing fragments were transferred directly to a small drop of demineralised water on a StarFrost adhesive microscopy slide (Cat. No.: 2510.1250, Hounisen Laboratorieudstyr, Skanderborg, Denmark) and allowed to dry, then outlined with a mini pap pen (ref.; 008877, Life Technologies). Primary antibodies diluted in IB with 0.1% Triton X-100 were applied for overnight incubation at 4°C. Myofibre typing fragments were then incubated with secondary antibodies and Hoechst 33342 diluted in IB for 2 h at 4°C, before the addition of prolong gold mounting medium (without DAPI) and a cover glass. Both the MTJ and myofibre typing fragments were washed two times with IB between all steps, followed by a final wash in TBS before mounting. Both fragments were left overnight at room temperature and stored at -20°C. This process was repeated several times per sample, to obtain 10 type I and 10 II myofibres for each of the 6 patients (120 total). In total 86 and 201 type I and II myofibres, respectively, were isolated and stained to reach this goal.

#### Single myofibres - RNAscope in situ hybridisation

RNAscope Multiplex Fluorescent Reagent Kit v2 (Cat. No: 323100. ACD Bio, Bio-Techne Ltd, Newark, USA) was used to visualise *COL22A1* transcripts in single myofibres from the *semitendinosus* muscle. Single fibre MTJ-fragments and myofibre typing fragments were obtained and the myofibre typing fragments used to determine the fibre type as described above (Fig. 6A,B). MTJ-fragments were transferred directly to a small drop of demineralized water on a StarFrost adhesive microscopy slide and allowed to dry. Afterwards, slides were dehydrated through an ethanol series (50%, 70%, 100% twice, for 5 min each at room temperature) and left in 100% ethanol at -20°C overnight. The following day the slides were dried for 5 min, and the MTJ- fragments outlined with a mini pap pen, before the addition of hydrogen peroxide (ACD Bio) for 10 min. After washing in distilled water, slides were protease-treated (Protease III, ACD Bio) for 15 min and washed twice in PBS. Probe hybridisation, with the *COL22A1* probe solution (ACD Bio), was carried out overnight at 40 °C in a HybEZ Slide Rack in a Humidity Control Tray (ACD Bio) with filter paper saturated with distilled water to prevent evaporation. The following day, the signal amplification was performed according to the manufacturer’s protocol. All incubations were at 40°C in a Humidity Control Tray (ACD Bio), followed by two 2 min washes using RNAscope wash buffer (ACD Bio). The first and second amplifiers were incubated for 30 min, the third was incubated for 15 min. The amplification was followed by 15 min incubation with horseradish peroxidase (ACD Bio), 30 min incubation with a tyramide dye fluorophore (Opal 620, Cat. No: FP1495001KT, Akoya Biosciences, Marlborough, USA) diluted 1:1000 in RNAscope TSA dilution buffer (ACD Bio), followed by 30 min incubation with a horseradish peroxidase blocker (ACD Bio). Here the protocol was modified by adding an immunofluorescence step, as described above (***single myofibres – immunofluorescence***), utilising a rabbit anti-dystrophin primary antibody (IgG, cat. no.: ab15277, Abcam, Cambridge, UK) together with a goat anti-rabbit DyLight 680 (Cat. No: 35568. Invitrogen) secondary antibody. Instead of Hoechst 33342, nuclei were labeled with RNAscope DAPI (ACD Bio) for 30 seconds directly after the final wash. Slides were mounted with Prolong Gold mounting medium and a cover glass.

### Image acquisition

Images of the myofibre typing fragments, longitudinal sections and cross-sectional sections were captured on an Olympus BX51 microscope with a 0.5x camera (Olympus DP71, Olympus Deutschland GmBH, Hamburg, Germany) using the Olympus cellSens Software (Version: Standard 1.14 (Build 14116), www.olympus-lifescience.com). Images of the myofibre typing fragments (Fig. 6B) and pig cross sections (Fig. 1A,B; Fig. S4) were captured using either a 10x (0.3 NA) or 40x (0.75 NA) objective and the human longitudinal sections were captured using either a 20x (0.5 NA) objective (Fig. 1C). High-resolution Z-stack images of immunofluorescence staining (Fig. 1D-E) of single myofiber MTJ fragments, together with RNAscope sample images (Fig. 4A), were captured using laser scanning confocal microscopy with a Zeiss LSM710 (Zeiss, Jena, Germany). An EC Plan-Neofluar 40x (1.3 NA) Oil DIC M27 objective (Software: Zeiss Zen Black 2012) or a Plan-Apochromat 20x (0.8 NA) Air M27 were used for either the high-resolution images or the RNAscope images, respectively. Nyquist sampling and a 1 Airy unit pinhole were used for all laser scanning confocal images. Hoechst 33342/DAPI, Alexa Fluor 594/OPAL620, and Alexa Fluor 680/DyLight 680 were excited by a 405 nm diode laser (30mW), a 561 nm solid-state laser (20mW) and a 633 nm HeNe laser (5mW), respectively. Images for 3D analysis of the MTJ fragments (Fig. 6C,D; Fig. S2), if properly aligned and undamaged, were captured using a spinning disk confocal microscope (CSU-X1, Yokogawa) mounted on a Zeiss CellObserver microscope, with an Orca Flash 4.0 LT sCMOS camera (Hamamatsu), using a Plan-Apochromat 20x (0.8 NA) objective (software: Zeiss Zen Blue 2012). Hoechst, Alexa Fluor 488, Alexa Fluor 568, and Alexa Fluor Phalloidin 680 were excited by a 405 nm diode laser, 488 nm argon laser, 561 HeNe laser and a 635 HeNe laser, respectively. Images were collected at a depth of 16 bits and the optical slice and Z-interval were adjusted to keep a constant voxel size of 0.325x0.325x0.35 µm for all images. Image parameters were kept the same for all images, except for laser intensities, which were adjusted to keep channel intensity profiles similar between fibres, to adjust for fluctuations in staining signal. Displayed images were cropped in Fiji (Schindelin et al., 2012) for presentation, and colour-blind friendly pseudo colours were applied to composite images. Additionally, using Fiji, a 3-pixel radii median filter was applied to all laser scanning confocal images for presentation.

### Image analysis

#### Single myofiber 3D analysis

In total 65 and 71 type I and II myofibres, respectively, were selected for imaging. For analysis of the nuclei 3D spatial organisation, the spinning disk confocal images of single myofibers (.czi) were converted to Imaris image files (.ims) and imported into Imaris (version 10.0, Oxford Instruments, https://imaris.oxinst.com/). The myofibre volumes were generated from the phalloidin staining channel using the Imaris surface segmentation function, and manually evaluated to ensure close overlap with the phalloidin and dystrophin staining. The 3D spatial coordinates of the nuclei were detected using the Imaris spot segmentation function. Excess spots were generated to ensure capture of all nuclei and were removed manually, preferentially leaving the centremost spot. Nuclei spots were then classified as either inside (myonuclei) or outside (nuclei of mononuclear cells) the myofibres, with a spot being classified as inside if located within the dystrophin staining.

Furthermore, we used vimentin as an additional marker to distinguish between inside and outside nuclei, if determination by dystrophin was ambiguous, as vimentin is widely known for perinuclear staining of mesenchymal cells (Ostrowska-Podhorodecka et al., 2022). Additionally, in previous work by our group, we observed by snRNA-seq that myonuclei isolated from human muscle-tendon tissue do not express vimentin (Fig. S3), further validating vimentin as a marker. If the Hoechst signal showed a major decrease in quality through the Z-stack, deconvolution of the Hoechst channel was performed in Imaris to improve nuclei segmentation. A full Imaris model display can be viewed in movie S2. Myofibres were excluded (a total of 8 myofibres) if proper segmentation could not be achieved. The 3D nuclei and surface objects were then analysed using an Imaris plugin (https://github.com/DBI-INFRA/Automated-muscle-fiber-analysis), created in Matlab (version 9.11, www.themathworks.com). In brief, the software imports the Imaris objects to Matlab and calculates the distance of nuclei spots to the myofibre tip, the distance of outside nuclei spots to the myofiber surface, the NN of inside nuclei spots, the MND of inside nuclei spots, and the outward-facing surface of MNDs (MNDS). All data are exported in an Excel sheet together with accompanying Matlab files. In total 55 type I and 57 type II myofibres were analysed, containing a total of 7969 and 6144 inside and outside nuclei, respectively. All MND and NN calculations for myonuclei bordering the open end of the myofiber were omitted from the analysis. For all included myofibres the MTJ-border was manually defined as the cessation of dystrophin folding, using maximum intensity Z-projections of the dystrophin channels in Fiji (Fig. 6D). The MTJ surface areas were calculated as the sum of all MNDS within the MTJ region.

#### RNAscope image analysis

Confocal images of RNAscope single myofibres (.CZI) were imported into Fiji, where measurements were made. Maximum intensity projection was used to collapse the three- dimensional Z-stacks into a two-dimensional image. A hand-drawn region-of-interest (ROI) surrounding the fibres was made using the dystrophin antibody staining as a marker, and the “Analyse Particles” function was used to create ROIs around each nucleus. Any nuclei ROI overlapping with or outside the myofibre dystrophin signal were excluded. *COL22A1* maximum intensity values were measured within the nuclei ROIs. Nuclei ROIs with a maximum intensity value above background levels were then manually evaluated for the expression of *COL22AI*, given by the presence of a clear fluorescent foci. In total 30 type I and 27 type II myofibres were included in the RNAscope *COL22A1* analysis.

### Statistical analysis

Plots and statistical comparisons were made in RStudio (RStudioTeam, 2022) running R version 4.2.0 (R Core Team, 2022). The following packages were used in R: Tidyverse 1.3.1, ggplot2 3.3.6, ggforce 0.3.3, ggrepel 0.9.1, readxl 1.4.0, xlsx 0.6.5, and ggpubr 0.4.0. A significance threshold of 0.05 was used. In preliminary experiments (data not shown) we observed 10.2 (mean, SD=2.9) and 4.1 (mean, SD=1.6) myonuclei at the MTJ vs a non-MTJ segment of single human myofibres, respectively, from which we estimated the need of 6 patients in the current study (power of 0.8). For all data points and statistics, patient median values (biological replicates) are used. For statistical assessments of MTJ cell numbers (Fig. 2C,D), the excess MTJ myonuclei (Fig. 2H), MTJ MND size (Fig. 3B), myonuclei MTJ-, adjacent-, and control zones (Fig. 3I-L), and quantification of *COL22A1*+ nuclei (Fig. 4B), a paired Wilcoxon signed rank test with continuity correction was performed. For the myonuclear number, MND, NN, and *COL22A1*+ nuclei profiles (Fig. 2E,F; Fig. 3C,D,F,G; Fig. 4C,D), a Friedman rank sum test was performed. Boxplots adhere to the Tukey method, showing interquartile ranges (IQR) with whiskers showing highest and lowest values within a 1.5 IQR distance from the 75^th^ and 25^th^ percentile, respectively.

## Acknowledgements

We thank Anja Jokipii-Utzon and Ann-Christina Ronnie Reimann for their technical assistance. We acknowledge the Core Facility for Integrated Microscopy, Faculty of Health and Medical Sciences, University of Copenhagen where the confocal imaging was performed. We acknowledge the Danish Bioimaging Infrastructure Image Analysis Core Facility for the support in the 3D model image analysis work presented. For providing the pig tissue, we are grateful to Stanislava Pankratova, Comparative Pediatrics and Nutrition, Department of Veterinary and Animal Sciences, Faculty of Health and Medical Sciences, University of Copenhagen.

## Competing interests

The authors declare no competing interests.

## Author contributions

Conceptualization: C.H., A.L.M., A.M., C.Y., M. Kjaer; Methodology: C.H., A.L.M., A.M., C.Y.; Formal Analysis: C.H., P.S., A.L.M., A.M., C.Y.; Investigation: C.H., A.M.; Resources: M. Koch, M.R.K.; Data Curation: C.H., A.M.; Visualization: C.H., A.M.; Writing – original draft: C.H., A.L.M.; Writing – review & editing: all authors; Supervision: A.L.M., C.C.Y., A.v.K.; Project administration: A.L.M.; Funding acquisition: M. Koch, M. Kjaer, A.L.M.

## Funding

Lundbeck Foundation (Grant ID: R344-2020-254 to A.L.M.), Novo Nordisk Fonden (Grant ID: NNF20OC0064829 to A.L.M.), Independent Research Fund Denmark (Grant ID: 10.46540/3101- 00063B to A.L.M.)

## Data availability

The custom software for the 3D model “Automated-muscle-fiber-analysis” can be found through the public github repository: https://github.com/DBI-INFRA/Automated-muscle-fiber-analysis

## Diversity and inclusion statement

In this study similar numbers of males and female patients were included.

## Notes

### Competing Interest Statement

The authors have declared no competing interest.

